# Oregano powder reduces *Streptococcus* and increases SCFA concentration in a mixed bacterial culture assay of chicken

**DOI:** 10.1101/625152

**Authors:** Benjamin W Bauer, Sheeana Gangadoo, Yadav Sharma Bajagai, Thi Thu Hao Van, Robert J Moore, Dragana Stanley

## Abstract

Food borne illnesses have a world-wide economic impact and industries are continuously developing technologies to reduce the spread of disease caused by microorganisms. Antimicrobial growth promoters (AGPs) have been used to decrease microbiological infections in animals and their potential transfer to humans. In recent years there has been a global trend to remove AGPs from animal feed in an attempt to reduce the spread of antimicrobial resistant genes into the human population. Phytobiotics, such as oregano powder, are one of the potential replacements for AGPs due to their well-established antimicrobial components. 16S rRNA gene amplicons were used to determine the effect of oregano powder (1% w/v) on the microbiota of mixed bacterial cell cultures, which were obtained from the ceca of traditionally grown meat chickens (broilers). Oregano powder had a mild effect on the microbial cell cultures increasing *Enterococcus faecium*, rearranging ratios of members in the genus *Lactobacillus* and significantly reducing the genus *Streptococcus* (*p*=1.6e^−3^). Beneficial short chain fatty acids (SCFA), acetic and butyric acid, were also significantly increased in oregano powder supplemented cultures. These results suggest that oregano powder at a concentration of 1 % (w/v) may have beneficial influences on mixed microbial communities and SCFA production.

## Introduction

Illness caused by the consumption of contaminated foods has a wide economic and public health impact worldwide [1]. In recent years, broiler producers have begun searching for alternatives to low dose sub-therapeutic AGPs that can maintain a healthy microbial gut flora without affecting the cost to poultry producers, the consumer or the environment. AGPs (e.g. avilamycin and zinc bacitracin) have been shown to reduce the abundance of some pathogenic groups of microorganisms, without having significant impacts on other members of the indigenous microbiota [2]. The ideal AGP alternatives would display similar influences over the microbial populations, while avoiding unforeseen problems with the health and performance of animals [3]. AGPs added to livestock and poultry feed have been shown to decrease zoonosis whilst improving animal health and performance [4, 5]. The occurrence of bacterial resistance in animal production facilities has led to the fear of the resistance genes spreading into the human population [4, 6–8], driving the European Union in banning AGPs, which has resulted in the global decrease of AGP usage [9–11]. However, reports have indicated that the sudden removal of AGPs may result in higher incidence of diseases in poultry farms, consequently increasing the use of therapeutic antibiotics [12, 13]. The efficiency of AGP alternatives need to be thoroughly investigated to ensure that the benefits to animals, consumers and producers are not lost. One such AGP alternative could be the use of oregano powder, replacing the system while maintaining similar benefits.

Oregano (*Origanum vulgare*) is a phytobiotic and is known to contain antimicrobial compounds such as carvacrol (CAR), thymol and their precursors, *p*-cymene and *γ*-terpinene, generally equating to 80 % of essential oil contents [14, 15]. CAR and thymol have consistently demonstrated antimicrobial properties at low doses [16–22], and reduced cell membrane potential which can lead to cell death [23]. This antimicrobial activity may have the potential to protect broilers from enteric pathogens and consumers from food borne diseases. Oregano powder has the potential to be used as a low dose subtherapeutic feed additive, preserving antibiotics for situations when high dose therapeutic treatments are required.

An assay utilising oregano, showed its superior influence with broiler weight gain and intestinal morphology when compared to avilamycin [24]. In another study, oregano, as a blend essential oil (25% thymol and 25% CAR), had a reduction in mortality, gut lesions and necrotic enteritis in broilers challenged with *Clostridium perfringens* [25], which cost the broiler industry billions of dollars per year [26]. Oregano essential oil has superior activity against pathogens compared to other herbs such as rosemary [27] and is a highly anti-oxidant [28], preservative [29–32] and anti-inflammatory substance [17, 19, 33].

In Australia, organic and layer chicken farmers restricted by the use of AGPs, are routinely adding oregano to poultry feed as an antibiotic alternative. However, the range of effects of dry rubbed oregano on pathogens and beneficial intestinal bacteria are unknown. Most of the previously mentioned studies were performed using classic culturing methods: plates, liquid cultures and disc diffusion assays. These methods targeted selected poultry pathogens in single cultures, showing that some bacterial species behave differently in the complexed microbial communities of the gastrointestinal tract [34]. Culturing a mixed bacterial culture could allow for the growth of unculturable strains due to cross-feeding of metabolic products and other complex community interactions (Lagier et al., 2012). The increased understanding of intestinal microbiota and its complex interactions have ensued different approaches towards microbiological media design, resulting in the successful cultivation of novel species, giving birth to a new area of microbiology - microbial culturomics [35].

Chickens have a short gastrointestinal feed retention time of approximately 3–4 h, and variable excreta microbiota due to periodic emptying of different gut sections [36]. Chicken caeca empty several times a day, continually sampling both ascending and descending microbiota through peristalsis and intestinal movement [37]. Thus, chicken caeca are considered as a reservoir of gut microbial diversity of all gastrointestinal sections. In this study we used modified and enriched LYHBHI microbiological media, capable of supporting a wide range of the caecal microbial community in the presence and absence of 1% oregano. The analysis of this complex bacterial community was performed using 16S rRNA gene sequencing. We present the first comprehensive study of the effects of oregano supplementation on caecal microbiota cultured in an in vitro model system.

## Materials and methods

### Animal ethics statement

The animal-based work in this study was approved and monitored by Central Queensland University Animal Ethics Committee under approval number A1409-318.

### LYHBHI Media Preparation and Enrichment

The LYHBHI media [Brain-heart infusion (37 g/L, BD), yeast extract (5g/L, Alfa Aesar), cellobiose (1g/L, BD), hemin (0.005 g/L, BD), L-cysteine (0.5 g/L, Alfa Aesar), resazurin sodium salt (5 mg/L, Alfa Aesar)], was initially diluted in 798 ml of water and autoclaved. After cooling down the media, 100 ml of filter sterilised bacterial ferment, 100 ml of chicken feed extract, 1 ml of vitamin and 1 ml of trace element mix was added to make up the volume up to 1 L of enhanced LYHBHI media. The media was then purged with (nitrogen) before use in an anaerobic chamber.

Bacterial ferment was prepared by culturing *Lactobacillus plantarum* (ATCC^®^ BAA-793™) and *Lactobacillus rhamnosus* (ATCC^®^ 53103™) in 60 ml of LYHBHI each in an aerobic incubator at 37°C, until stationery phase. The cultures were centrifuged and 50 ml of supernatant from both cultures was mixed and filter sterilised to enrich the LYHBHI media.

Feed extract was prepared by grinding on full power (1500 W, Nutri Ninja Auto iQ Duo, SharkNinja, USA) 100 g of chick starter crumble (Red Hen Chick premium micro starter crumbles antibiotic and hormone free, Lauke Mills, Daveystone SA, Australia) for 3 min. One litre of water was added to the powdered feed and blended again for an additional 3 min. The mixture was then autoclaved, allowed to cool overnight then centrifuged (3220 rcf, 5 min) in 50 ml falcon tubes. The clear supernatant was then pooled, filter sterilised and 100 ml of filtrate added to lukewarm LYHBHI.

Vitamin mix was prepared by dissolving a capsule of vitamin mix (Multivitamins and Minerals, Cenovis, Aust) and a capsule of vitamin K2 (Caruso’s Natural Health, Australia) in 10 ml of water separately and filter sterilising 1 ml of resulting solutions was added to 1 L of LYHBHI media. Trace elements mix (Youngevity, California, USA) was filter sterilised (1 ml) and added to the enriched media. Both vitamin and trace mineral mix concentrations in final 1 L of enriched LYHBHI is provided in Supplementary File 1, Table 1.

**Table 1.** The concentration of CAR in cultures shown in mmol/L. CAR concentrations were determined by GCMS analysis.

| **Condition** | **$\eta$** | **1A** | **1B** | **2A** | **2B** | **3A** | **3B** |
| --- | --- | --- | --- | --- | --- | --- | --- |
| Cont | 3 | 0 | 0 | 0 | 0 | 0 | 0 |
| Oreg | 3 | 0.2 | 0.21 | 0.2 | 0.19 | 0.19 | 0.19 |

### Caecum Starter Cultures

The complete caeca from three roosters (6 months old) were donated to the laboratory by a local organic heritage breeder (Rockhampton, QLD, AUS). The caeca were delivered frozen and in anaerobic conditions. Before culturing caeca were placed in the anaerobic work station (A35, Whitley, Shipley, UK) at 37°C using mixed gas (CO_2_ 10%, H_2_ 10%, N_2_ 80% BOC, QLD, Australia) until they were completely thawed. The caecal contents were aseptically emptied into Erlenmeyer flasks containing 50 ml of enhanced LYHBHI media inside the anaerobic work station. To ensure good transfer of mucosa-associated bacteria, the whole caeca were then opened laterally its contents and tissue transferred into the prepared Erlenmeyer flasks. Cotton stoppers were placed in the mouth of each flask to allow for gas exchange. The cultures from all three rooster’s caeca were incubated for 2 hrs in the anaerobic chamber at 37 °C on the orbital shaker (89 rpm). After 2 hrs the cultures were aliquoted into Cryo-tubes with 700 μl culture and 300 μl sterile glycerol in each tube and stored at −80°C.

### Oregano powder preparation

The dried aerial parts of oregano were used to make the powder. The oregano was processed by blending (100 g, 1.5 min/max, 1500 W, Nutri Ninja Auto iQ Duo) to reduce particle size. That powder was then processed in a Planetary Ball Mill Machine (speed no. 5, 2 hrs, 40 g*each run, Changsha Yonglekang Equipment, China). A representative sample of each stage of processing was kept in a cool dark place. The oregano was then placed in an electric sieve machine (Changsha Yonglekang Equipment, China) to collect material that passed through the 75μm, sieve.

### Inoculation of Caecal Microbial Culture with Oregano

One aliquot of caecum starter culture of each of the three roosters was gradually thawed by slowly increasing temperature. Enhanced LYHBHI media was purged using nitrogen in an anaerobic workstation. The media was aliquoted (20 ml) into twelve 50 ml Erlenmeyer flasks. Oregano powder was added to each of the six treatment flasks at 1% w/v. The flasks were plugged with cotton stoppers to allow gas exchange and incubated at 37 °C under a CO_2_ 10%, H_2_ 10%, N_2_ 80% gas mix in an orbital shaker (100 rpm) in the anaerobic workstation. In a separate experiment we established that the cultures reached stationary phase before 24 hours. The cultures were incubated for 24 hrs and then centrifuged at 1300 rpm for 10 min. The supernatant was removed, and stored at −80 °C for short chain fatty acid (SCFA) analysis. The pellet was processed for 16S rRNA amplicon sequencing.

### DNA Extraction

The pellet obtained from centrifuged cultures was used to extract total DNA for 16S rRNA sequencing. The pellets were resuspended, in the remaining media, using a vortex and transferred into tubes containing 0.2 g of glass beads (0.1 mm, diameter) and 0.7 ml of lysis buffer (500 mM NaCl, 50 mM EDTA, 50 mM TrisHCl (pH8), 4% SDS). Samples were homogenised at maximum speed for 5 min using a beadbeater (Mini-Beadbeater, Biospec products). The samples were then incubated at 75 °C for 15 min vortex at 5 min intervals. Samples were then centrifuged (16,000 rcf, 5 min) and 0.4 ml of the supernatant was transferred to a 1.5 ml tubes that contained 500 μl of binding buffer (5 M Gu-HCl, 30% isopropanol). The samples were vortexed and then centrifuged (16,000 rcf, 1 min). All of the supernatant was transferred into a DNA spin column with collection tube (Enzymax LLC, Cat# EZC101, Kentucky, US). The spin column was centrifuged (8,000 rcf, 1min) and the contents of collection tube discarded. The spin column was then washed twice with 800 μl of wash buffer (10 mM Tris-HCl, 80% ethanol (pH=7.5) centrifuging at 8,000 rcf for 1 min. The spin columns were dried by centrifugation (8,000 rcf, 1min) and placed in new collection tubes. and eluted with 50 μl of elution buffer (10 mM Tris-HCl). The DNAs were analysed in a NanoDrop spectrophotometer to determine concentration and quality.

### 16S rRNA Sequencing

The primers to amplify the V3-V4 region of 16S rRNA genes were: forward ACTCCTACGGGAGGCAGCAG and reverse GGACTACHVGGGTWTCTAAT. The primers contained barcodes, spacers and Illumina sequencing linkers were designed and used as previously described. The sequencing was conducted on an Illumina MiSeq instrument using 2×300 bp paired-end sequencing.

Microbial communities were initially analysed using Quantitative Insights Into Microbial Ecology (QIIME v.1.9.1) [38]. Paired end sequences were joined using Fastq-Join algorithm and with no mismatches allowed within the region of overlap. Phred quality threshold was minimum 20. OTUs were picked at 97% similarity using Uclust [39] and inspected for chimeric sequences using Pintail [40]. All taxonomic assignments were performed in QIIME against the GreenGenes database and QIIME default arguments [41]. Unifrac matrix was calculated in QIIME and on the rarefied OTU table. OTUs with less than 0.01% abundance were removed. Statistical analysis including Spearman correlations, alpha and beta diversity were done on Hellinger transformed data [42, 43]. Significantly differentially abundant taxa were analysed using negative binomial distribution based DeSeq2 [44] method for differential analysis of sequencing count data. DESeq is performed on raw sequence counts.

Further data exploration was done in Calypso [45]. Sequencing data is publicly available on MG-RAST database under accession number MglX (pending).

### Metabolite Extractions for SCFA Analysis

Culture supernatant samples were thawed on ice and 1 ml of each SCFA sample was transferred into a capped 2 ml tube containing 1 ml of 70% ethanol. The sample was then filtered through a 0.45μm cellulose syringe filter into a 2 ml vial. Standards of acetic acid, butyric acid, propanoic, valeric acid, isobutyric acid were prepared in 70% ethanol at concentrations of 1 ppm, 10 ppm, 50 ppm, 100 ppm, 500 ppm, 700 ppm, 1000 ppm. CAR was prepared in the same manner with concentrations of 10 ppm, 20 ppm, 40 ppm, 80 ppm and 100 ppm.

### GC-MS systems and methods

The samples and standards were then placed in a GC-MS (GC-MS-QP2010 Ultra fitted with an AOC-20s Shimadzu auto sampler and a Shimadzu AOC-20i auto injector) with a polar column (Agilent J&W GC, 30m, 0.250 diam (mm), film 0.25 (μm) temperature limits form 40°C to 260°C). SCFA was determined by injecting 1 μl of sample at 250 °C with helium (1.97 ml/min) as the carrier gas with a 5.0 split injection mode. Pressure was maintained at 143.3 kPa and helium flow of 103.4 ml/min. The mass spectrometer operated in the electron ionization mode at 0.2kV, the source temperature was 220 °C with scan mode between 33 to 150*m/z*.

CAR was detected by injecting 1 μl of sample at 250 °C with helium (1.7 ml/min, 5.0) as the carrier gas with a 50.0 split injection mode. Pressure was maintained at 161.1 kPa and helium flow of 89.7 ml/min. The mass spectrometer operated in the electron ionization mode at 0.2 kV, the source temperature was 230 °C with scan mode between 33 to 280 *m/z*. Total program time was 5.88 min. The National Institute of Standards and Technology (NIST) library was used to identify and match peaks.

## Results

### Microbiota supported by enhanced LYHBHI

The caecal phyla supported by the media included (in order of abundance) Firmicutes, Proteobacteria, Actinobacteria, Bacteroidetes, Tenericutes and Spirochaetes. These phyla were represented by a total of 50 genera, 23 of which are uncultured. The cultured genera included: *Arthrobacter*, *Bacteroides*, *Bifidobacterium*, *Clostridium*, *Collinsella*, *Coprobacillus*, *Dehalobacterium*, *Dorea*, *Enterococcus*, *Epulopiscium*, *Erwinia*, *Eubacterium*, *Faecalibacterium*, *Lactobacillus*, *Lactococcus*, *Ochrobactrum*, *Oscillospira*, *Pediococcus*, *Proteus*, *RFN20*, *Ruminococcus*, *Slackia*, *Sphaerochaeta*, *Streptococcus*, *Sutterella* and *Trichococcus* with unclassified genera from families Streptococcaceae, Enterobacteriaceae and Lactobacillales among the most abundant. Figure 1 shows the 20 most abundant genera supported by enhanced LYHBHI media. The high number of unclassified genera members in the original caeca is a consequence of the impact of non-industrialised traditional housing and a very different environmental conditions [46] such as fully ranging outdoor roosting birds, high live plant and insect food content, sharing yard with other poultry species and exposure to wild birds and animals.

**Figure 1:**
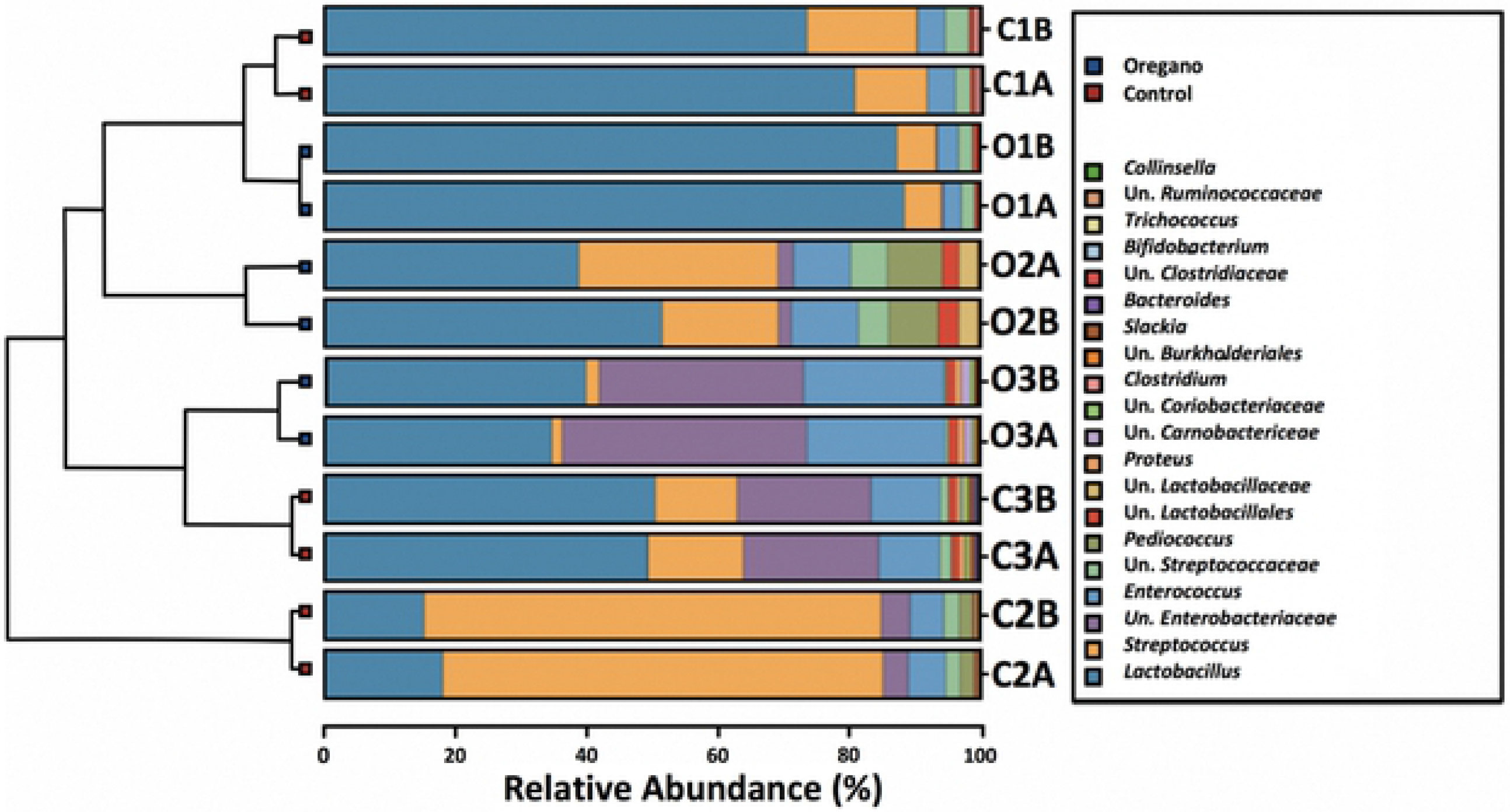
Hierarchical sample clustering bar-chart showing 20 most abundant genera. Sample names start with C or O (control or oregano), number (rooster 1, 2 or 3) and replicate code (A or B for 2 replicates).

### Alpha and beta diversity

Alpha diversity was not affected by the addition of 1% oregano based on Richness (*p*=0.96), Evenness (*p* = 0.54), Shannon index (*p* = 0.58) or Simpson index (*p* = 0.43). Beta diversity indicators did not suggest major community perturbations by oregano with OTU level permutational multivariate analysis of variance (PERMANOVA) (Bray-Curtis) showing no significant oregano effect (*p* = 0.166). However, the microbiota biological donor had substantial influence on microbial community with rooster impact *p* = 3.3E^−4^ (Figure 2 A, B).

**Figure 2:**
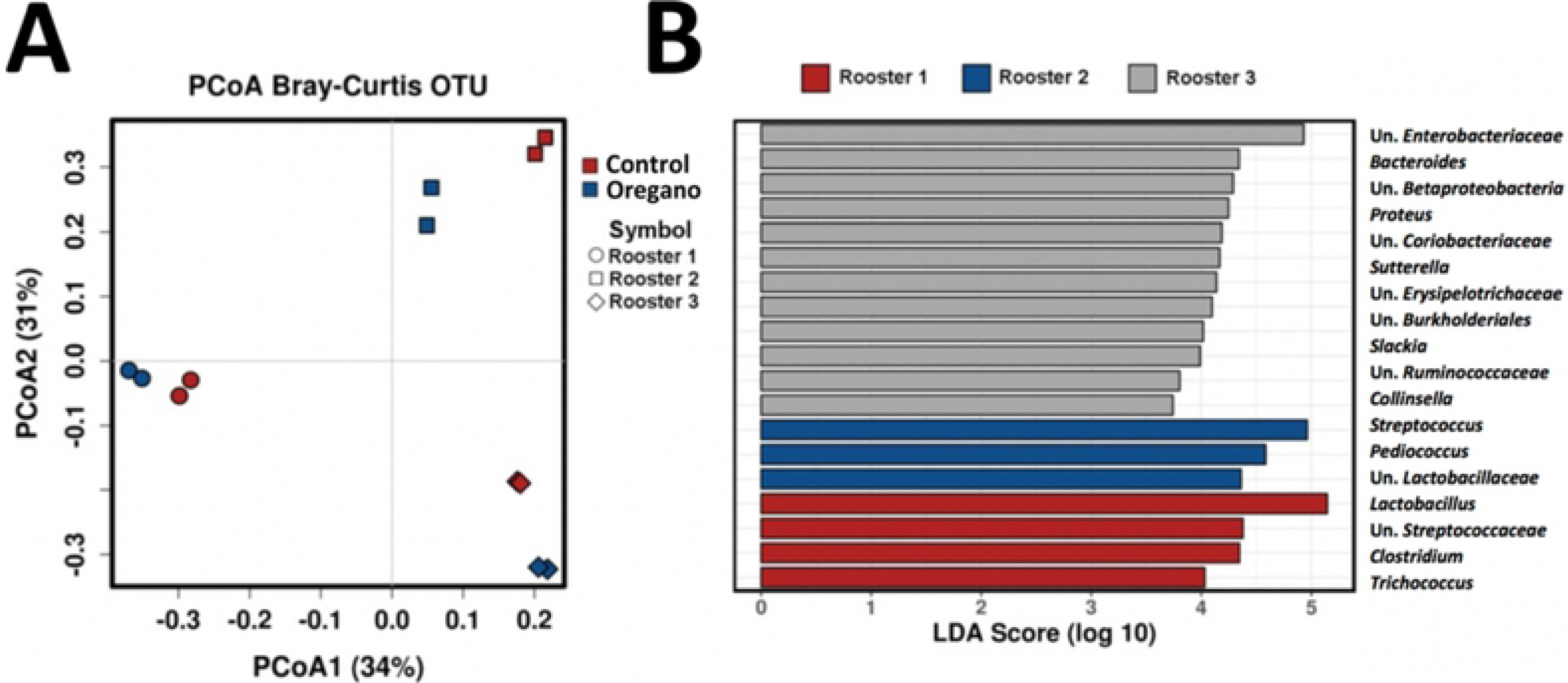
High effect of biological microbiota donor. Panel A shows Bray-Curtis PCoA plot separating samples by the rooster donor rather than oregano; panel B shows linear discriminant analysis (LDA) effect size method showing genera likely responsible for difference between the rooster’s microbiotas.

We used DeSeq2 negative binomial analysis to identify differentially abundant features. The biggest effect was seen in: *Streptococcus* (*p* = 0.016), was 60 % reduced and an unclassified genus of Lactobacillales (*p* = 0.013) 2.4 fold increased in oregano treatment group (Figure 3 A-D). 96 OTUs were significantly different in abundance between the treatments. Among a number of those belonging to unclassified genera, there were also 62 differentially abundant OTUs from *Lactobacillus*, 15 from *Streptococcus* and 5 from *Enterococcus* genera. Both *Streptococcus* and *Enterococcus* OTUs were consistently changed in the same direction, all 15 *Streptococcus* OTUs were reduced (Figure 3 B, C) while all 5 *Enterococcus* OTUs were increased by oregano. Blastn analysis identified the most abundant *Streptococcus* OTU (Figure 3 B) to be highly similar to *Streptococcus gallolyticus* strain St5, with 100 % sequence identity and an e-value=0. Oregano treatment reduced this OTU in rooster 2 caecal community, where it was the most abundant, from 55.7 % to 17.4 % in one and from 51.1 % to 9.7 % reads in another replicate. Most of the other *Streptococcus* OTUs aligned with uncultured *Streptococcus*. The *Enterococcus* OTUs were either 100 % identical to *Enterococcus faecium* strain HBUAS54015 or to uncultured bacterial clone database targets.

**Figure 3:**
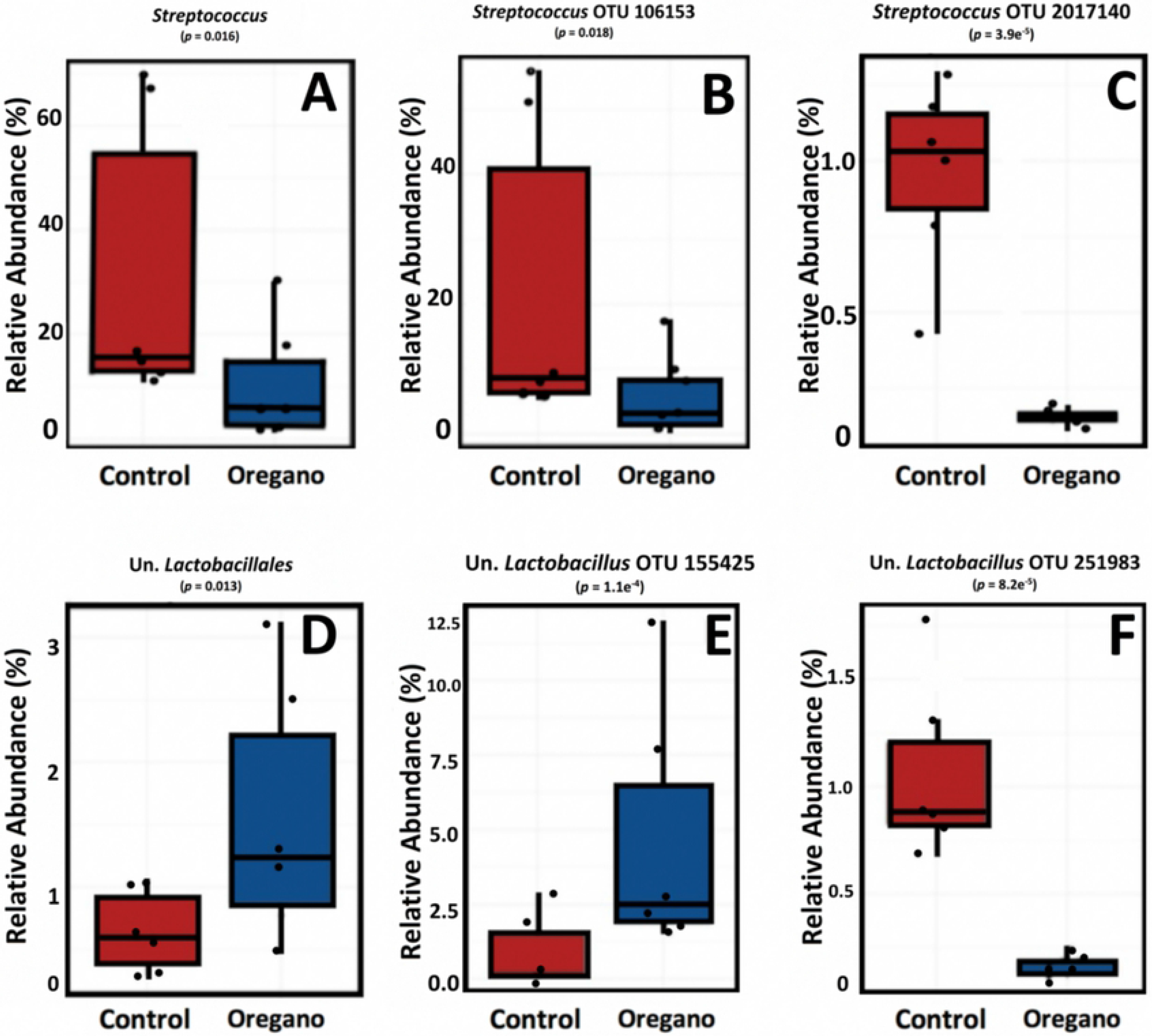
DeSeq2 differentially abundant taxa. Table with all significant OTUs is provided in Supplementary File 1, Table 2.

The *Lactobacillus* genus was not significantly changed (*p* = 0.46), however, 62 *Lactobacillus* OTUs were significantly altered by oregano treatment, some significantly increased (Figure 3 E) and others reduced (Figure 3 F). Based on blastn analysis OTUs 100 % similar to *Lactobacillus crispatus* (shown in Figure 3 E) or *Lactobacillus ingluviei* were significantly increased and other OTUs similar to uncultured *Lactobacillus* bacterial clone database targets (shown in Figure 3 F) were reduced by oregano treatment. All significant OTUs with their closest blast hit IDs are given in Supplementary File 1, Table S2.

### Oregano influenced SCFA production

The main SCFA produced in cultures were acetate, butyrate, proponate, valerate and isobutyrate. There were significantly increased amounts of acetate (*p* = 6.6e^−4^) and butyrate (*p* = 6.3e^−3^) in oregano treatment groups (Figure 4). Overall on average oregano groups produced 61% more of the total 5 SCFA analysed (*p =* 0.022).

**Figure 4:**
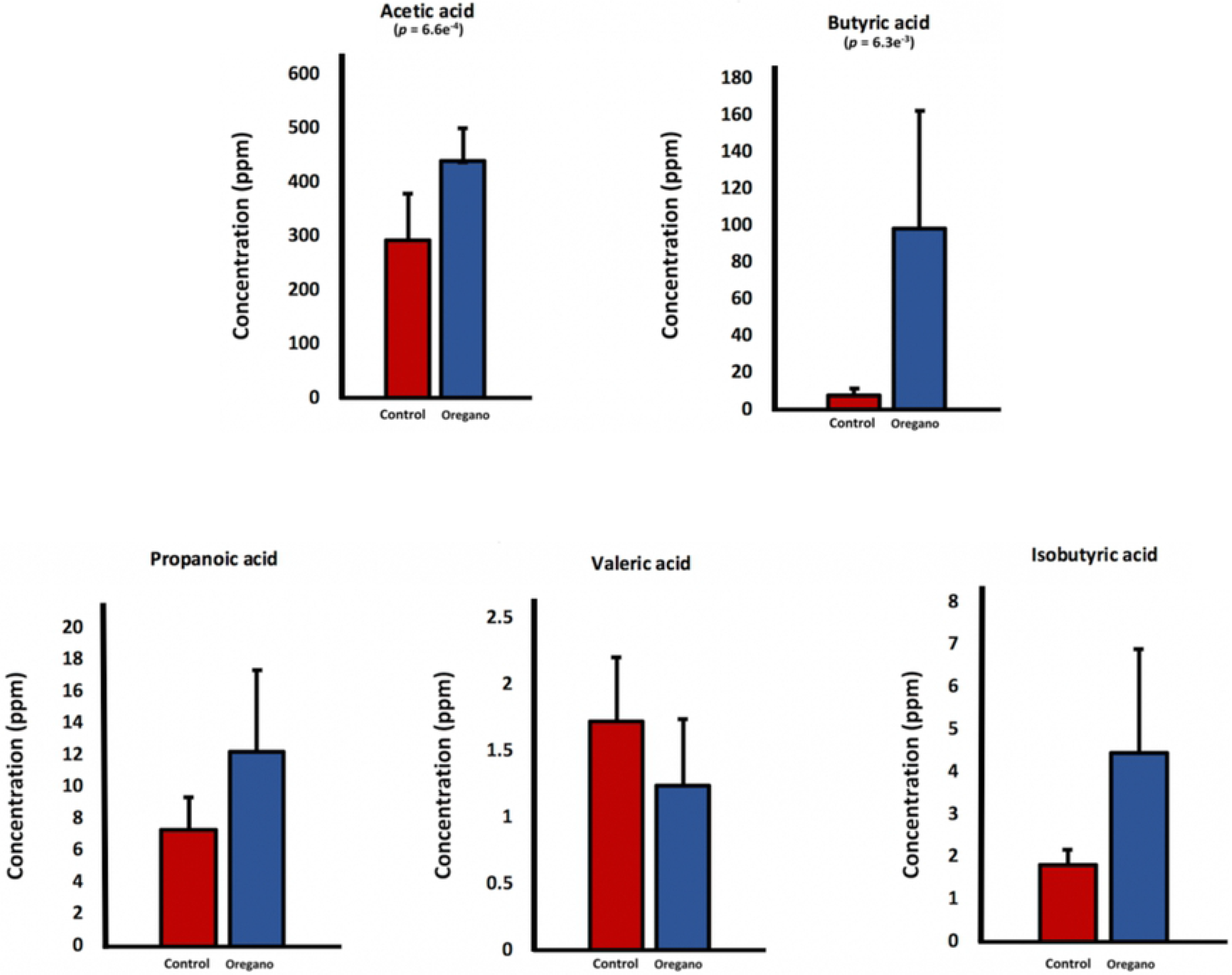
Concentrations of SCFA in control and oregano supplemented cultures Figure 4: Concentrations of SCFA in control and oregano supplemented cultures. One-way Anova was used to determine significance. There were two SCFA significantly increased, which were AA (*p* = 6.6*e*^−4^) and BA (*p* = 6.3*e*^−3^)

### SCFA production correlated with some microbial taxa

We used PERMANOVA (on Bray-Curtis matrix) to evaluate if concentrations of produced SCFA, namely acetate, butyrate, isobutyrate, proponate and valerate had significant interactions with microbial communities. Only acetate (*p* = 6.6e^−4^) and butyrate (*p* = 6.3e^−3^) showed significant interaction with microbiota. Since we did not supplement SCFA to the culture, it is not possible to distinguish if oregano increased the abundance of high SCFA-producing microbiota, or if oregano stimulated existing bacteria to make more SCFA. It is also impossible to infer if microbiota influenced increase of SCFA or increased SCFA influenced microbiota structure. Spearman’s correlation test showed a number of significant interactions between concentrations of AA (Supplementary File 1, Table S3, Figure 5A) and of BA (Supplementary File 1, Table S4, Figure 5B) with bacterial genera.

**Figure 5:**
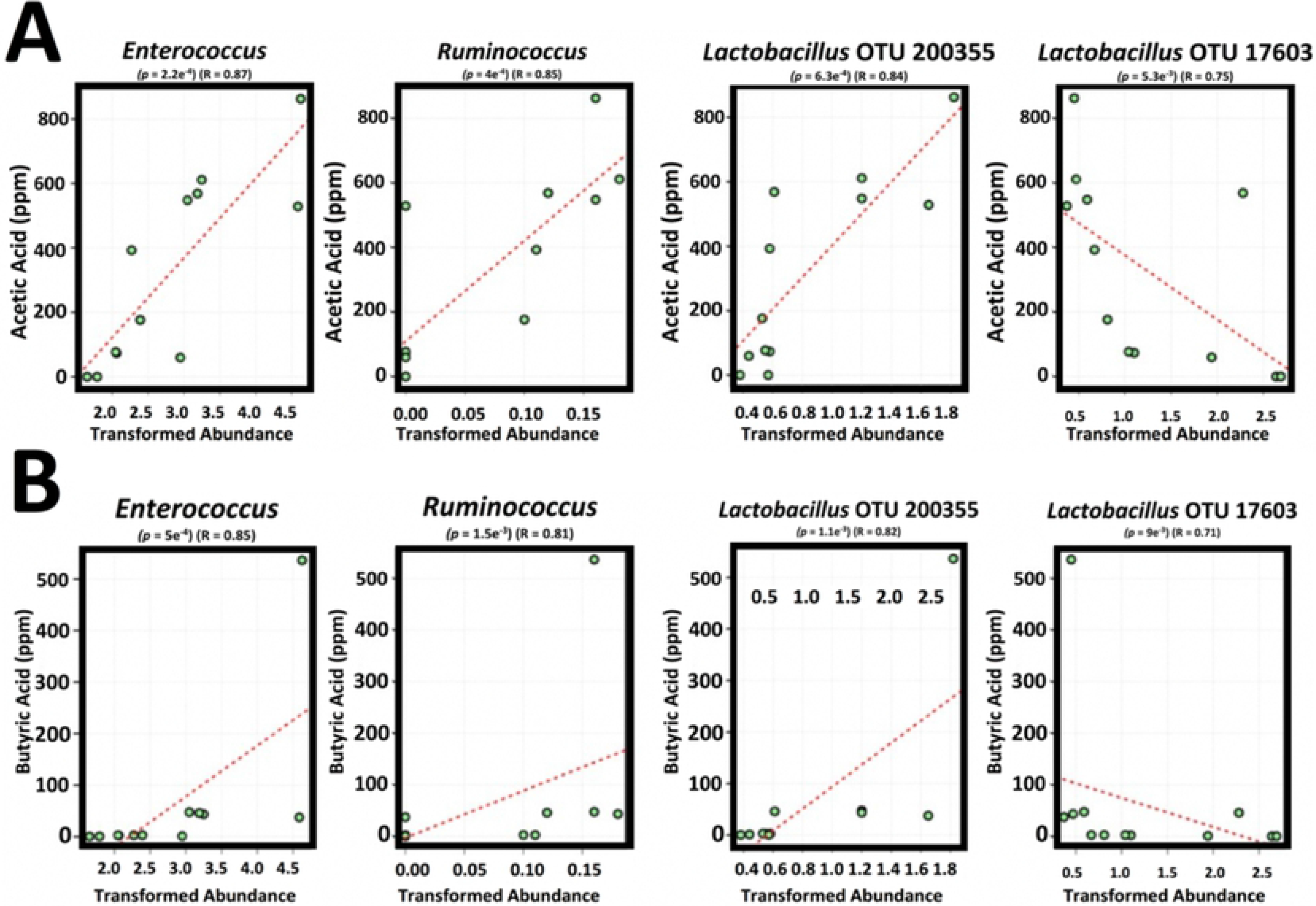
SCFA: AA and BA spearman correlations of concentration and abundance.

Acetate significantly (p < 0.05) positively correlated with *Enterococcus*, *Ruminococcus*, *Proteus*, *Coprobacillus*, *Bacteroides*, *Sutterella*, *Bifidobacterium* and *Collinsella* and negatively correlated with *Clostridium* while butyric had very similar interaction profile to acetate, with positive interactions with *Enterococcus*, *Ruminococcus*, *Proteus*, *Coprobacillus*, *Sutterella* and *Bacteroide*s and negative correlation with *Clostridium*. Among the top 20 most abundant OTUs the significant correlations with acetate and butyrate were almost identical, with *Lactobacillus* OTU 200355 positively correlated with both acetate (*p* = 6.3E−4, R=0.84) and with butyrate (*p* = 1.1e^−3^, R = 0.82) while another *Lactobacillus* OTU 17603 showed significant negative correlations with both acetate (*p* = 5.3e^−3^, R = −0.75) and BA (p = 0.009, R = −0.71). The other significantly correlated OTUs, especially the most abundant ones, showed the same pattern, with very similar response to acetate and butyrate (Figure 6). Other SCFAs also had a number of highly significant correlations, however, did not have significant influence on microbial community structure (Figure 6).

**Figure 6:**
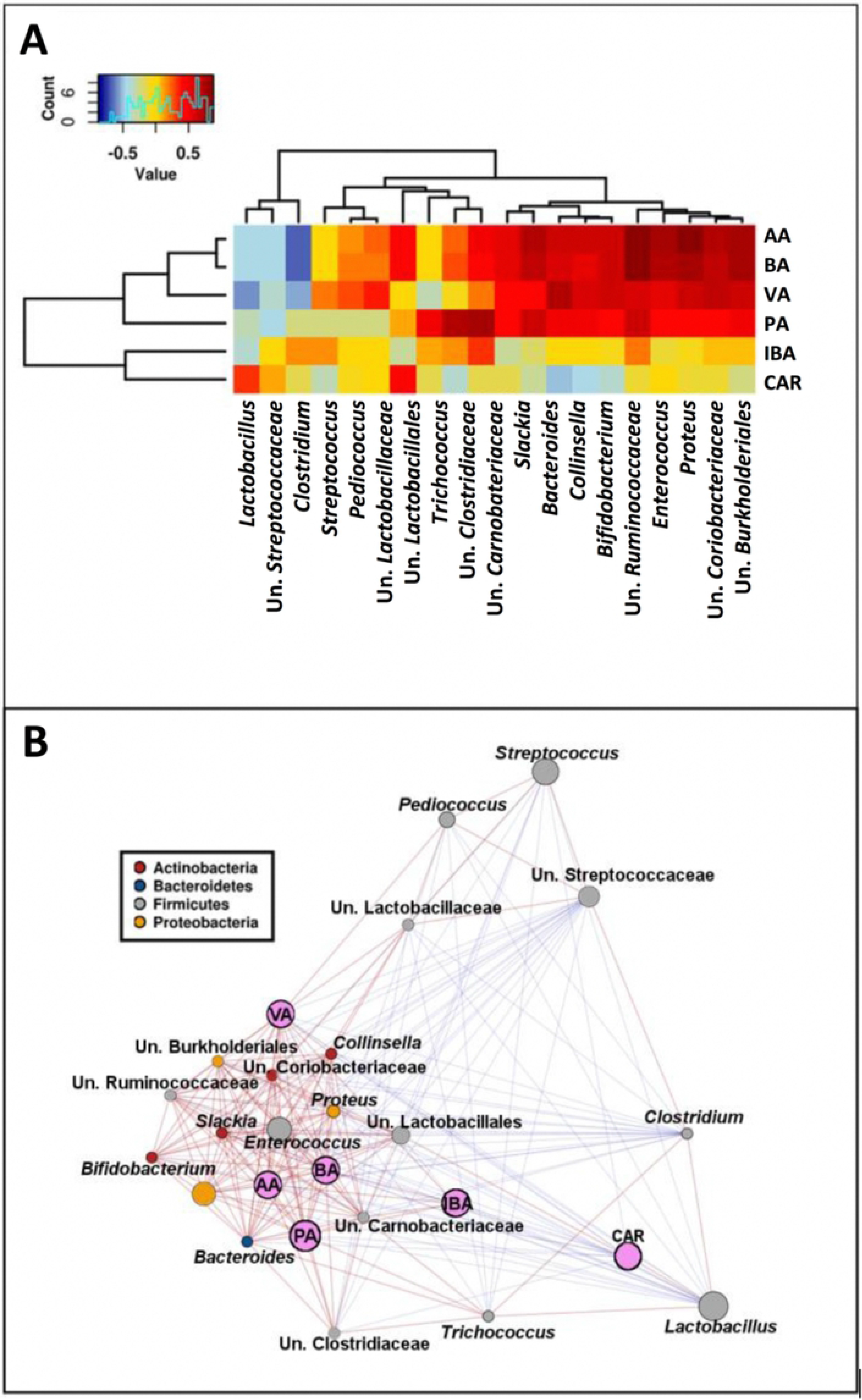
Heatmap (A) and a network (B) showing 20 most abundant genera relationship with SCFA and CAR. Both figures are based on Spearman correlation.

### CAR and microbial community

As expected CAR was not detected in the control group (Table 1), and based on PERMANOVA performed on Bray-Curtis matrix at an OTU level, CAR had marginally significant influence on microbial communities (*p* = 0.044). There were no genera significantly correlated with concentration of carvacrol in the microbial cultures and overall effect of acetate and butyrate appeared to overpower borderline significance of carvacrol influence. OTUs that correlated with CAR concentrations are provided in Supplementary File 1 Table S5.

## Discussion

The gastrointestinal tract (GIT) microbiota has been extensively studied because of the significant role it plays in the health and performance of animals, including poultry [47–49]. Earlier studies on GIT microbiota were mostly, invitro, culture-based techniques. However, traditional methods are unable to cultivate the majority of the GIT microbiota and thus had limitations to unravel the complexity of this ecosystem. It has been estimated that only 10 – 50 % of chicken caecal bacteria can be cultured [37]. In this assay, both control and treatment cultures had a wide range of unknown bacteria, this could be an indication that the enhanced LYHBHI media may be beneficial in culturing fastidious bacteria. The co-culture of diverse caecal microbial metabolic products can cross-feed bacteria facilitating their growth[50]. This fact could also explain the presence of unidentified bacteria within the cultures.

Oregano, at 1 % concentration, did not have a destructive effect on the caecal microbiota, in fact, its effects were largely overpowered by the biological variation originating in the caeca of the different roosters. Instead, oregano selectively targeted certain bacterial groups, especially reducing *Streptococcus*, increasing *Enterococcus* and re-arranged ratios of *Lactobacillus* species, without affecting the total *Lactobacillus* abundance.

Reduction of *Streptococcus* (*p* = 1.6e^−3^) species by 1 % oregano media supplementation was not surprising as CAR has been shown in numerous assays, to inhibit the growth of various *Streptococcus* species, such as *S. pyogenes*, *S. mitis*, *S. mutans*, *S. sanguis*, *S. milleri* and *S*. *pneumoniae* [51–53]. However, the heterogeneity of oregano, particularly the concentrations of other antimicrobial phytochemicals, such as thymol, *p*-cymene and *y*-terpinene, may have contributed to the reduction of *Streptococcus*, alongside variations in the complexed microbiota. In the cultures, the most abundant *Streptococcus* OTU identified with 100 % sequence identity across the amplified region was *Streptococcus gallolyticus*, it was reduced in one rooster’s caecal community, where it was the most abundant, from 55.7 % to 17.4 % and from 51.1 % to 9.7 % reads in another replicate. *Streptococcus gallolyticus* is known to cause septic bacteraemia in preterm neonates [54] as it has been strongly associated with intestinal damage and colorectal cancer [55, 56] as well as colonising colorectal tumours [57], endocarditis [58] and other associated health issues. This indicated possible benefit of oregano for individuals genetically predisposed to colorectal cancer. In poultry production this could protect against zoonosis of pathogenic *Streptococcus* strains to consumers.

The Enterococcus OTUs promoted by oregano were related (100 % sequence id) to *Enterococcus faecium*, a well-known intestinal coloniser. *Enterococcus faecium* strains can be either alpha-haemolytic or non-haemolytic, extremely pathogenic or probiotic. Pathogenic strains represent a major health issue as a Vancomycin-resistant *E. faecium*, referred to as VRE [59] Other strains are marketed as probiotics improving intestinal health [60] and immunity [61]. In chickens probiotic strains of *E. faecium* show a positive effect in birds infected with *Salmonella* sp. [62]. An *E. faecium* probiotic strain was shown to promote growth performance, improve intestinal morphology, and improve the caecal microflora in *Escherichia coli* challenged broilers [63].

Rearrangements in representatives of the *Lactobacillus* genus can have either positive or negative influence on the bird performance. In human research, *Lactobacillus* are becoming well known as obesogenic [64] despite being beneficial on other health fronts. On the other hand, other *Lactobacillus* strains were shown as weight loss promoting [65]. The same discrepancy was reported in chicken feed efficiency, with some *Lactobacillus* strains positively correlated and others negatively correlated with weight gain or productivity [66]. Moreover, it was shown that the outcome of *Lactobacillus*, either as a probiotic or in weight loss and productivity, is strictly strain level specific. Fak and Backhed have shown that two different strains of *L. reuteri* had opposite effect on weight loss in mice [67]. Thus, changes in the abundance of different *Lactobacillus* species may promote or reduce more beneficial *Lactobacillus* strains and influence bird heath and productivity.

Despite the influence of oregano on *Lactobacillus* sp. and *E. faecium* being strain specific, restricting definite inference of beneficial effects using the 16S-based methodology, the SCFAs were quantitatively measured and shown to be increased by 1 % of oregano. The health promoting effects are well documented. Butyrate has been shown to be the preferred energy source for colonic cells, and increases the production of tight junction proteins [68], improves maintenance and structure of villi [12], and reduces inflammation [69, 70], which may improve the growth and performance of broilers experiencing stress [71]. Acetate has shown to be a potent antimicrobial [72] against *Streptococcus* and *Salmonella* [73] and additionally can cross-feed butyrate production. SCFA production was 61 % higher in treatment groups, in live birds these increases would translate to healthier microbial communities, intestinal morphology and immune systems of broilers that receive oregano powder as a feed supplement. The observed increased levels of acetate and butyrate in treatment groups suggests that in vivo application of oregano powder at 1%, could benefit the host by providing energy to intestinal cells, anti-inflammation and antimicrobial activities resulting in assistance with competitive exclusion.

## Supporting information

SupplementaryFile1.pdf

## Acknowledgments

We wish to acknowledge Jason Bell and help he provided in all aspects of High-Performance Computing. We also thank Corine Ting, Vicky Carroll and Charmaine Elder for technical support also heritage chicken breeder Kerrilyn Salmond for providing the samples that were used for this assay’s cultures. This study, including scholarship for BB, was funded by Poultry CRC established by Australian Government.

## References

1. Gandhi M, Chikindas ML. Listeria: A foodborne pathogen that knows how to survive. International Journal of Food Microbiology. 2007;113(1):1–15.

2. Crisol-Martinez E, et al. Understanding the mechanisms of zinc bacitracin and avilamycin on animal production: linking gut microbiota and growth performance in chickens. Applied Microbiology and Biotechnology. 2017;101(11):4547–4559.

3. Lozupone CA, et al. Diversity, stability and resilience of the human gut microbiota. Nature. 2012;489(7415):220–230.

4. Emborg HD, et al. The effect of discontinuing the use of antimicrobial growth promoters on the productivity in the Danish broiler production. Preventive Veterinary Medicine. 2001;50(1-2):53–70.

5. Casewell M, et al. The European ban on growth-promoting antibiotics and emerging consequences for human and animal health. Journal of Antimicrobial Chemotherapy. 2003;52(2):159–61.

6. Turnidge J. The Use of Antibiotics in Food-Producing Animals: Antibiotic-Resistant Bacteria in Animals and Humans. F.a.F. Commonwealth Department of Agrictulture, Editor. 1999, Australian Government:Canberra ACT 2601.

7. Jensen HH, Hayes DJ. Impact of Denmark’s ban on antimicrobials for growth promotion. Curr Opin Microbiol. 2014;19:30–36.

8. Turnidge J. Antibiotic use in animals--prejudices, perceptions and realities. Journal of Antimicrobial Chemotherapy. 2004;53(1):26–7.

9. Betancourt L, et al. Effect of Origanum chemotypes on broiler intestinal bacteria. Poultary Science. 2014;93(10):2526–35.

10. Karimi A, et al. Effects of level and source of oregano leaf in starter diets for broiler chicks. The Journal of Applied Poultry Research. 2010;19(2):137–145.

11. Windisch W, et al. Use of phytogenic products as feed additives for swine and poultry. Journal of Animal Science. 2008;86(14 Suppl):E140–8.

12. Onrust L, et al. Steering Endogenous Butyrate Production in the Intestinal Tract of Broilers as a Tool to Improve Gut Health. Frontiers in Veterinary Science. 2015;2:75.

13. Phillips I, et al. Does the use of antibiotics in food animals pose a risk to human health? A critical review of published data. Journal of Antimicrobial Chemotherapy. 2004;53(1):28–52.

14. Brenes A, Roura E. Essential oils in poultry nutrition: Main effects and modes of action. Animal Feed Science and Technology. 2010;158(1-2):1–14.

15. Giannenas I, et al. Effect of dietary supplementation with oregano essential oil on performance of broilers after experimental infection with eimeria tenella. Archives of Animal Nutrition. 2003;57(2):99–106.

16. Boskabady MH, Jalali S. Effect of carvacrol on tracheal responsiveness, inflammatory mediators, total and differential WBC count in blood of sensitized guinea pigs. Experimental Biology and Medicine. 2013;238(2):200–8.

17. Arigesavan K, Sudhandiran G. Carvacrol exhibits anti-oxidant and anti-inflammatory effects against 1, 2-dimethyl hydrazine plus dextran sodium sulfate induced inflammation associated carcinogenicity in the colon of Fischer 344 rats. Biochemical and Biophysical Research Communications. 2015;461(2):314–20.

18. Lima Mda S, et al. Anti-inflammatory effects of carvacrol: evidence for a key role of interleukin-10. European Journal of Pharmacology. 2013;699(1-3):112–7.

19. Silva FV, et al. Anti-inflammatory and anti-ulcer activities of carvacrol, a monoterpene present in the essential oil of oregano. Journal of Medicinal Food. 2012;15(11):984–91.

20. Dagli Gul AS, et al. The effects of oral carvacrol treatment against H2O2 induced injury on isolated pancreas islet cells of rats. Islets. 2013;5(4):149–55.

21. De Vincenzi M, et al. Constituents of aromatic plants: carvacrol. Fitoterapia. 2004;75(7-8):801–4.

22. Sivropoulou A, Papanikolaou E, Nikolaou C, Kokkini S, Lanaras T, Arsenakis M. Antimicrobial and Cytotoxic Activities of Origanum Essential Oils. Journal of Agriculture and Food Chemistry. 1996;44:4.

23. Ultee A, Bennik MHJ, Moezelaar R. The phenolic hydroxyl group of carvacrol is essential for action against the food-borne pathogen Bacillus cereus. Applied and Environmental Microbiology. 2002;68(4):1561–1568.

24. Peng QY, et al. Effects of dietary supplementation with oregano essential oil on growth performance, carcass traits and jejunal morphology in broiler chickens. Animal Feed Science and Technology. 2016;214:148–153.

25. Yin D, et al. Supplemental thymol and carvacrol increases ileum Lactobacillus population and reduces effect of necrotic enteritis caused by Clostridium perfringes in chickens. Scientific Reports. 2017;7(1):7334.

26. Van Immerseel F, et al. Rethinking our understanding of the pathogenesis of necrotic enteritis in chickens. Trends in Microbiology. 2009; 17(1):32–36.

27. Mathlouthi N, et al. Use of rosemary, oregano, and a commercial blend of essential oils in broiler chickens: in vitro antimicrobial activities and effects on growth performance. Journal of Animal Science. 2012;90(3):813–23.

28. Young JF, et al. Ascorbic acid, alpha-tocopherol, and oregano supplements reduce stress-induced deterioration of chicken meat quality. Poultry Science. 2003;82(8):1343–51.

29. Bajpai VK, Baek KH, Kang SC. Control of Salmonella in foods by using essential oils: A review. Food Research International. 2012;45(2):722–734.

30. Hyldgaard M, Mygind T, Meyer RI. Essential oils in food preservation: mode of action, synergies, and interactions with food matrix components. Frontiers in Microbiology, 2012;3:12.

31. Lagouri V, et al. Composition and antioxidant activity of essential oils from Oregano plants grown wild in Greece. Zeitschrift fur Lebensmittel-Untersuchung und-Forschung. 1993;197(1):20–23.

32. Olmedo R, Nepote V, Grosso NR. Antioxidant activity of fractions from oregano essential oils obtained by molecular distillation. Food Chemistry. 2014;156:212–9.

33. Ocana-Fuentes A, et al. Supercritical fluid extraction of oregano (Origanum vulgare) essentials oils: anti-inflammatory properties based on cytokine response on THP-1 macrophages. Food and Chemical Toxicology. 2010;48(6):1568–75.

34. Nicholson JK, et al. Host-gut microbiota metabolic interactions. Science. 2012;336(6086):1262–7.

35. Lagier JC, et al. Microbial culturomics: paradigm shift in the human gut microbiome study. Clinical Microbiological and Infection. 2012;18(12):1185–93.

36. Svihus B. Function of the digestive system. Journal of Applied Poultry Research. 2014;23(2):306–314.

37. Stanley D, Hughes RJ, Moore RJ. Microbiota of the chicken gastrointestinal tract: influence on health, productivity and disease. Applied Microbiology and Biotechnology, 2014;98(10):4301–10.

38. Caporaso JG, et al. QIIME allows analysis of high-throughput community sequencing data. Nature Methods. 2010;7(5):335–336.

39. Edgar RC. Search and clustering orders of magnitude faster than BLAST. Bioinformatics. 2010;26(19):2460–2461.

40. Ashelford KE, et al. At least 1 in 20 16S rRNA sequence records currently held in public repositories is estimated to contain substantial anomalies. Applied and Environmental Microbiology. 2005;71(12):7724–7736.

41. DeSantis TZ, et al. Greengenes, a chimera-checked 16S rRNA gene database and workbench compatible with ARB. Applied and Environmental Microbiology. 2006;72(7):5069–5072.

42. Legendre P, Gallagher ED. Ecologically meaningful transformations for ordination of species data. Oecologia. 2001;129(2):271–280.

43. Legendre P, De Caceres M. Beta diversity as the variance of community data: dissimilarity coefficients and partitioning. Ecology Letters, 2013;16(8):951–963.

44. Love MI, Huber W, Anders S. Moderated estimation of fold change and dispersion for RNA-seq data with DESeq2. Genome Biology. 2014;15(12).

45. Zakrzewski M, et al. Calypso: a user-friendly web-server for mining and visualizing microbiome-environment interactions. Bioinformatics. 2017;33(5):782–783.

46. Kers JG, et al. Host and environmental factors affecting the intestinal microbiota in chickens. Frontiers in Microbiology. 2018;9:235.

47. Ahlman H Nilsson O. The gut as the largets endocrine organ in the body. Annals of Oncology. 2001;2:6.

48. Deplancke B, Gaskins HR. Microbial modulation of innate defense: goblet cells and the intestinal mucus layer. The American Journal of Clinical Nutrition. 2001;73(6):11.

49. Ursell LK, et al. Defining the human microbiome. Nutrition Reviews. 2012;70 Suppl 1:S38–44.

50. Seth EC, Taga ME. Nutrient cross-feeding in the microbial world. Frontiers in Microbiology. 2014;5.

51. Magi G, Marini E, Facinelli B. Antimicrobial activity of essential oils and carvacrol, and synergy of carvacrol and erythromycin, against clinical, erythromycin-resistant Group A Streptococci. Frontiers in Microbiology. 2015;6:165.

52. Lakis ZM, Nicorescu D, Vulturescu V, Udeanu DI. The Antimicrobial Activity of Thymus Vulgaris and Origanum Syriacum Essential Oils on Staphylococcus Aureus, Streptococcus Pneumoniae and Candida Albicans. Farmacia. 2012;60(6):857–865.

53. Didry N, Dubreuil L, Pinkas M. Activity of thymol, carvacrol, cinnamaldehyde and eugenol on oral bacteria. Pharmaceutica Acta Helvetiae. 1994;69:4.

54. Floret N, et al. A cluster of bloodstream infections caused by Streptococcus gallolyticus subspecies pasteurianus that involved 5 preterm neonates in a university hospital during a 2-month period. Infection Control and Hospital Epidemiology. 2010;31(2):194–196.

55. Boleij A, et al. Clinical Importance of Streptococcus gallolyticus Infection Among Colorectal Cancer Patients: Systematic Review and Meta-analysis. Clinical Infectious Diseases. 2011;53(9):870–878.

56. Abdulamir AS, et al. Investigation into the controversial association of Streptococcus gallolyticus with colorectal cancer and adenoma. BMC Cancer. 2009;9:12.

57. Abdulamir AS, Hafidh RR, Abu Bakar F. Molecular detection, quantification, and isolation of Streptococcus gallolyticus bacteria colonizing colorectal tumors: inflammation-driven potential of carcinogenesis via IL-1, COX-2, and IL-8. Molecular Cancer. 2010;9:18.

58. Rusniok C, et al. Genome Sequence of Streptococcus gallolyticus: Insights into Its Adaptation to the Bovine Rumen and Its Ability To Cause Endocarditis. Journal of Bacteriology. 2010;192(8):2266–2276.

59. Jordens JZ, Bates J, Griffiths DT. Faecal carriage and nosocomial spread of vancomycin-resistant Enterococcus faecium. Journal of Antimicrobial Chemotherapy. 1994;34(4):515–28.

60. Scharek L, et al. Influence of a probiotic Enterococcus faecium strain on development of the immune system of sows and piglets. Vet Immunol Immunopathol. 2005;105(1-2):151–61.

61. Benyacoub J, et al., Supplementation of food with Enterococcus faecium (SF68) stimulates immune functions in young dogs. The Journal of Nutrition. 2003;133(4):1158–62.

62. Carina Audisio M, Oliver G, Apella MC. Protective effect of Enterococcus faecium J96, a potential probiotic strain, on chicks infected with Salmonella Pullorum. Journal of Food Protection. 2000;63(10):1333–7.

63. Cao GT, et al. Effects of a probiotic, Enterococcus faecium, on growth performance, intestinal morphology, immune response, and cecal microflora in broiler chickens challenged with Escherichia coli K88. Poultry Science. 2013;92(11):2949–55.

64. Million M, et al. Obesity-associated gut microbiota is enriched in Lactobacillus reuteri and depleted in Bifidobacterium animalis and Methanobrevibacter smithii. International Journal of Obesity. 2012;36(6):817–825.

65. Sanchez M, et al. Effect of Lactobacillus rhamnosus CGMCC1.3724 supplementation on weight loss and maintenance in obese men and women. British Journal of Nutrition. 2014;111(8):1507–19.

66. Stanley D, et al. Bacteria within the Gastrointestinal Tract Microbiota Correlated with Improved Growth and Feed Conversion: Challenges Presented for the Identification of Performance Enhancing Probiotic Bacteria. Frontiers in Microbiology. 2016;7.

67. Fak F, Backhed F. Lactobacillus reuteri prevents diet-induced obesity, but not atherosclerosis, in a strain dependent fashion in Apoe−/− mice. PLoS One. 2012;7(10):e46837.

68. Peng L, et al. Butyrate enhances the intestinal barrier by facilitating tight junction assembly via activation of AMP-activated protein kinase in Caco-2 cell monolayers. The Journal of Nutrition. 2009;139(9):1619–25.

69. Leeson S, et al. Effect of butyric acid on the performance and carcass yield of broiler chickens. Poultry Science. 2005;84(9):1418–22.

70. Beh BK, et al. Anti-obesity and anti-inflammatory effects of synthetic acetic acid vinegar and Nipa vinegar on high-fat-diet-induced obese mice. Sci Rep. 2017;7(1):2045–2322.

71. Zhang WH, et al. Sodium butyrate maintains growth performance by regulating the immune response in broiler chickens. British Poultry Science. 2011;52(3):292–301.

72. Ryssel H, et al. The antimicrobial effect of acetic acid--an alternative to common local antiseptics? Burns. 2009;35(5):695–700.

73. Andreatti RL, et al. Use of Anaerobic Cecal Microflora, Lactose and Acetic Acid for the Protection of Broiler Chicks against Experimental Infection with Salmonella typhimurium and Salmonella enteritidis. Brazilian Journal of Microbiology. 2000;31(2):107–112.

